# Endothelial Kallikrein-Related Peptidase 8 Promotes Diabetic Nephropathy through a LIFR dependent mechanism

**DOI:** 10.1101/2025.07.01.662673

**Authors:** Jiankui Du, Mingyang Li, Ying Jiang, Zhengshan Tang, Danhong Xu, Hongling Yin, Jihong Yuan, Xiaoyan Zhu, Xin Ni

## Abstract

**Background:** Diabetic nephropathy (DN) is the primary microvascular complication of diabetes mellitus; however, the exact pathways in endothelial cells (ECs) linked to DN progress remain unclear. Tissue kallikrein-related peptidases (KLKs) participate in pathophysiological processes in ECs. We aimed to explore the roles of endothelial KLKs in DN and define the underlying mechanisms.

**Methods and results:** KLK8 was the most highly upregulated member of KLKs in renal tissues in streptozotocin (STZ)-induced diabetic mice and cultured glomerular ECs (GECs) upon high glucose (HG) treatment. Both global (Klk8^-/-^) and endothelial Klk8 knockout (Klk8^ΔEC^) mice displayed improved albuminuria, mesangial matrix expansion and glomerulosclerosis caused by STZ compared to Klk8^f/f^ mice. Single-cell RNA seq (scRNA-seq) showed that many pathways associated with DN in ECs, mesangial cells (MCs) and tubule cells were reversed in Klk8^ΔEC^ mice. Endothelial-to-mesenchymal transition (EndMT) was extensively improved in Klk8^ΔEC^ mice, and by KLK8 siRNA in cultured GECs upon HG. Using proteome and other biochemical approaches, we revealed that KLK8 cleaved syndecan-4(SDC4), which contributed to loss of glycocalyx integrity in GECs in cultured cells and animal diabetic models. Furthermore, scRNA-seq showed that Lifr was one of the key genes linked to the disease progressed in ECs and MCs and regulated by endothelial Klk8. LIFR signaling contributed to HG-induced GEC dysfunction and MC activation, which was associated with endothelial Klk8. Knockdown of Lifr by lentivirus-Lifr shRNA ameliorated hallmark features of DN and improved EndMT and Sdc4 expression glomeruli of diabetic mice. LIFR was upregulated in GECs and MCs in DN patients. Circulatory levels of LIF, KLK8 and soluble SDC4 were increased in patients with DN, and KLK8 level was positively correlated with LIF, soluble SDC4 and creatinine levels.

**Conclusion:** Endothelial KLK8 promotes GECs dysfunction and abnormal crosstalk with MCs through a LIFR-dependent signaling in DN, which immediately highlights therapeutic targets for DN.

**Graphical Abstract:** 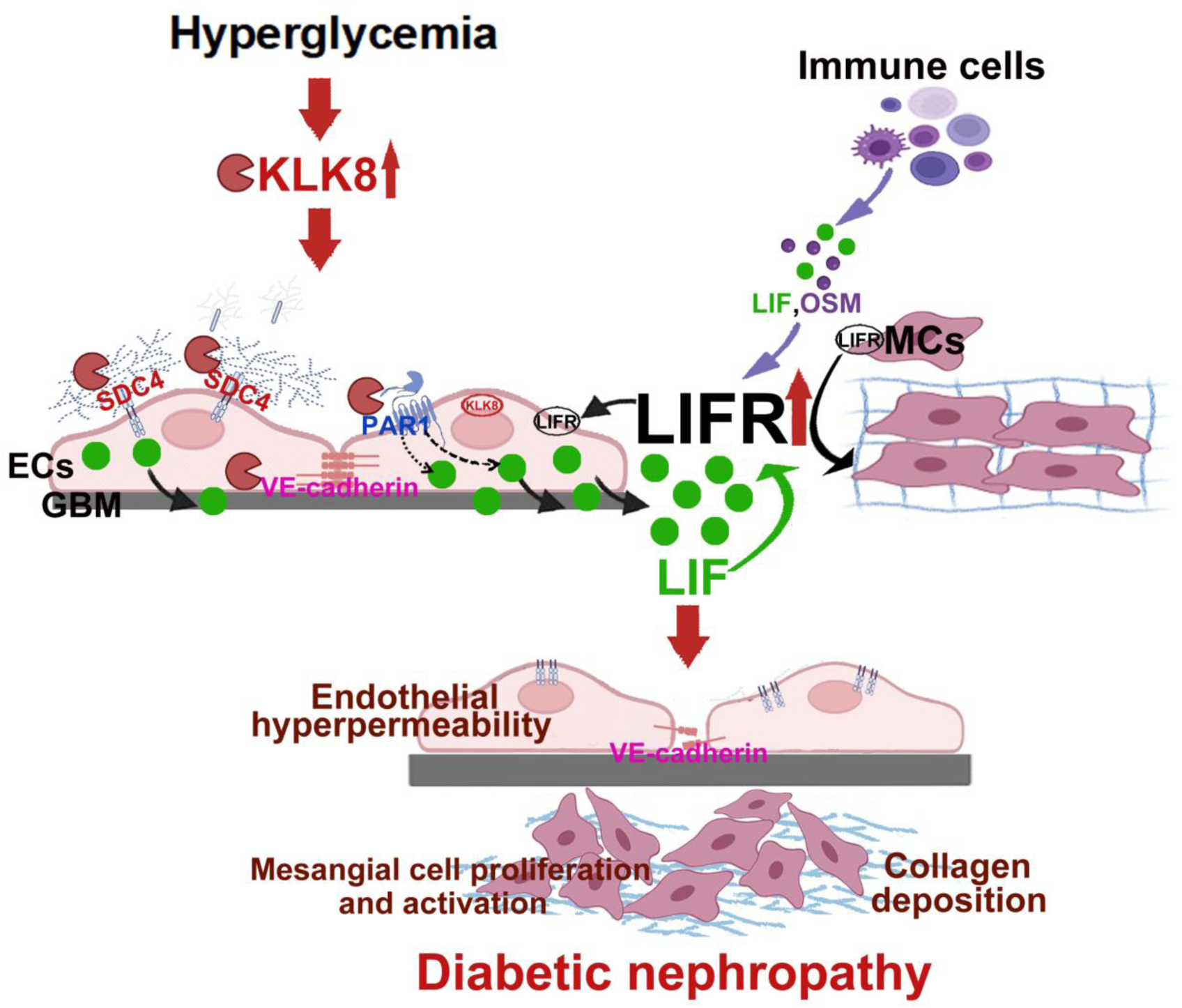

## Introduction

Diabetic nephropathy (DN) is one of the most frequent and serious complications of diabetes, and the leading cause of end-stage renal disease worldwide. About 40% of diabetic patients develop DN, and many diabetic patients continue to progress to end-stage kidney disease despite intensive glycemic and blood pressure management^1^. Thus, elucidating the molecular mechanisms intrinsic to the development of DN is essential for the formulation of effective intervention strategies.

DN is the primary microvascular complication of diabetes mellitus; with abnormal vascular hemodynamics and disturbed vascular permeability leading to clinical important features, such as microalbuminuria. Emerging evidence has demonstrated that some endogenous factors produced within the endothelial cells (ECs) are involved in DN progress, with being activated or inhibited in diabetes; therefore, they could potentially be used alongside traditional treatments. For instance, the important roles of dysregulation of vasoactive factors, such as vascular endothelial growth factor A (VEGFA), angiopoietins-1(Ang-1) and-2 (Ang-2), and endothelial nitric oxide synthase (eNOS) in DN pathogenesis have been reported^2–4^. Recently, endothelial leucine-rich α-2 glycoprotein-1(LRG1) is found to be involved in DN progress through Tgf-β signaling^5^. However, endothelial factors linked to DN development and progress remain largely unknown.

The tissue kallikrein-related peptidase (KLK) family consists of at least 15 genes in humans located on chromosome 19q13.3-13.4. KLKs share homology in gene structure, protein sequence, and tertiary structure, and are secreted into most biological fluids and are expressed in nearly all healthy tissues^6^. They can mediate diverse physiological functions and pathological processes through kinin-dependent and independent mechanisms. In ECs, KLKs contribute to various functions and processes including vascular homeostasis^7^, angiogenesis, inflammation^8^, thrombosis^9^, etc. For instance, KLKs regulate blood pressure and vascular tone through the activation of the kallikrein-kinin system. KLK1 and KLK3 are involved in angiogenesis by degrading extracellular matrix (ECM) proteins and facilitating EC migration and capillary formation^10,11^. As far as DN is concerned, some studies indicate that KLK1 plays protective roles in DN development. Global Klk1 deficiency worsens albuminuria and glomerulosclerosis in streptozotocin (STZ)-induced diabetic mice, while administration of exogenous kallikrein prevents and ameliorates DN in STZ-induced diabetic mice and obese type 2 diabetic mice^12,13^. Kallikrein activity is reduced in plasma in diabetic patients, which is associated with reduced kidney function^14^. More recently, Kobayashi et al. have proposed that circulatory KLK8 level is a risk factor for DN development and progress in humans^15^. Thus, it becomes of great interest in understanding the roles of KLKs in DN development and progress.

The aims of the present study were to explore the roles of KLKs in DN progress and define the underlying mechanisms. At first, we demonstrated that renal KLK8 is the most highly upregulated member of KLKs in ECs and diabetic animal models. We then revealed that endothelial KLK8 plays a crucial role in DN through a LIFR-dependent mechanism, and circulatory KLK8 level is associated with DN progress in diabetic patients by using multiple approaches including mouse genetics, single-cell transcriptome, and proteome, biochemical approaches. Our study reveals a novel key endothelial factor in DN pathogenesis and immediately highlights therapeutic targets for DN.

## Methods

### Human Samples

Blood samples were obtained from patients with DN (n = 20) and healthy controls (n = 20). DN was clinically diagnosed according to the guidelines of DN^16^, ie, UACR > 300 mg/g and eGFR < 90 mL/min/1.73 m^2^. The samples of healthy controls were obtained from the people who underwent routine health check-ups at Xiangya Hospital Central South University. Renal biopsy tissues were obtained from patients with DN (n = 7) and those with minimally changed nephrotic syndrome (MCNS) (n = 7). They were diagnosed as DN according to the above guidelines and morphology confirmation. The clinical characteristics of the patients with DN enrolled in this study were shown in Table S1& S2. Ethics approval was obtained from the Ethics Committee of Medical Research of Xiangya Hospital Central South University (No.2023091177). Informed consent was obtained from all participants, and the study was performed according to the declaration of Helsinki.

### Animal models

All experimental steps were supported by the Ethical Committee of Medical Research of Xiangya Hospital Central South University (Changsha, China). Animal studies were performed in accordance with the Guide for the Care and Use of Laboratory Animals published by the NIH (NIH publication No. 85-23, revised 1996). All laboratory mice in this study were maintained in a pathogen-free facility at the Animal Research Center of Navy Medical University.

Global Klk8 knockout (Klk8^-/-^) mice generation and identification were performed according to the previously described^17^. Briefly, Klk8-floxed mice were mated with EIIa-Cre transgenic mice (The Jackson Laboratory). Deletion of the Klk8 gene was verified by PCR of genomic DNA using (5’-GGACGTTGGAGTCACAGC-3’) and (5’-CCCAGGAGCAGAAGAGTG-3’) primers. Klk8^flox/flox^/EIIa-Cre(+) mice (KLK8^-/-^), along with age-matched Klk8^flox/flox^/EIIa-Cre(-) littermates (Klk8^f/f^) as controls, were used to investigate the impacts of Klk8 deficiency. Since single-cell sequencing data showed that Tie2 mRNA is mainly expressed in endothelial cells (GSE244475), we generated endothelial-specific Klk8 knockout mice by crossing Tie2-Cre transgenic mice (The Jackson Laboratory) with Klk8-floxed mice. Klk8^flox/flox^/Tie2-Cre^(+)^ mice, ie, Klk8^ΔEC^ mice, along with age-matched Klk8^flox/flox^; Tie2-Cre^(-)^ littermates (Klk8^f/f^) as controls, were used to investigate the impacts of endothelial-specific Klk8 deficiency. Adult mice with the same genotype were assigned to 1:1 allocation ratio to control and diabetic groups. Diabetes was induced by the intraperitoneal injection of 100 mg/kg streptozotocin (STZ, Sigma Aldrich, St. Louis, MO) in citrate buffer (pH 4.5) for 2 consecutive days as described previously^17^. Mice with non-fasting glucose levels of ≥ 250 mg/dL on day 4 after injection were considered diabetic. The mice were maintained until 16 weeks after STZ injection, then GFR was determined. Mice were sacrificed with 2.5% isoflurane via inhalation and cervical dislocation where appropriate after collecting urine, and blood and renal tissues were then collected. The samples were snap-frozen in liquid nitrogen and stored at -80°C for further analysis. Additionally, some renal tissues were fixed in paraformaldehyde (4%).

To investigate the role of LIFR in the progression of DN, adult C57/BL/6 mice were purchased from the Shanghai SLAC Laboratory Animal Co. (Shanghai, China), and diabetes was induced by STZ. Diabetic mice were randomly divided into three groups: one vehicle control group and two diabetic groups. One week after STZ injection, mice in one diabetic group were injected with lentivirus carrying Lifr short hairpin RNA (shRNA), while mice in another diabetic group were injected with lentivirus carrying control shRNA. The constructs of lentivirus GV493 (pFU-GW-016) vector carrying Lifr shRNA (Lv-Lifr shRNA) were constructed by Shanghai GenePharma Co (Shanghai, China). The sequences of shRNA targeting mouse Lifr gene were GGACATCAATTCAACAGTTGT and those of negative control were TTCTCCGAACGTGTCACGT. The amount of lentivirus was 1×10^12^ transducing units. The mice were sacrificed after collection of urine samples at 16^th^ week after STZ injection. Blood and renal tissues were then collected, snap-frozen in liquid nitrogen, and stored in -80°C for further analysis. Additionally, some renal tissues were fixed in paraformaldehyde (4%).

### Single-cell RNA-Seq (scRNA-seq)

The scRNA-seq was conducted by Novogene Bioinformatics Technology Co., Ltd (Tianjing, China). Briefly, mouse kidneys were collected and chilled in ice-cold PBS, followed by dissection of the cortex tissue. The tissues were initially minced and then digested with a digestion solution at 37°C for 30-40 mins. Single-cell suspensions were prepared by filtering through a 40-μm cell strainer and then washed three times with PBS at 4°C. Cell viability was determined by trypan blue staining with a TC20 automated cell counter (Bio-rad, Hercules, CA), yielding a viability rate exceeding 80%. The concentration of the single-cell suspension was adjusted to 700-1200 cells/μL. The cells of each sample were then processed using the MobiCube® High Throughput Single Cell 3’RNA-Seq Kit (v2.1) following manufacturer’s instruction in Novogene Bioinformatics Technology.

### Single-Cell Data Analysis

We used Fastp to perform basic quality control and statistics on the raw reads. The raw scRNA-seq data included Read1, Read2, and the i7 index read. Read1 contained the sequence of the Barcode and UMI (unique molecular identifiers), while Read2 was the sequence of the cDNA fragment. The raw reads of Read1 and Read2 was qualified for subsequent analysis. The raw data was analyzed by MobiVision v3.2 (MobiDrop) with default parameters. After processing, the raw reads were aligned to the reference genome, and a gene-cell matrix was generated. This matrix served as input data for further analysis with Seurat.

Cell types for each cluster were identified based on the similarity of enriched genes in each cluster compared to marker genes of known cell types, as described in prior studies^18,19^. Marker genes for each cell type were provided in supplemental File.1. Differential expression analysis between groups for each cell type was conducted using the Seurat function ’FindMarkers’ to identify significantly differentially expressed genes (DEGs). Enrichment analysis and Gene Set Enrichment Analysis (GSEA) were performed using the R package clusterProfiler (v4.14.6). Up-regulated and down-regulated genes were extracted from the DEG list for enrichment analysis, selecting the top 1000 genes with p_val < 0.05 and the largest absolute avg_log2FC values. All genes with p_val< 0.05 in the DEG list were used for GSEA analysis. Enrichment terms with p_value < 0.05 were considered significant enrichment pathways. Cell-Cell interaction analysis was performed using the R-package CellChat (v2.1.2). Data from the "CellChatDB.mouse" database were extracted to create CellChat objects for Control/Klk8^f/f^, STZ/Klk8^f/f^ and STZ/Klk8^ΔEC^. The receptor-ligand pair information in the "CellChatDB.mouse" database included Secreted Signaling, ECM-Receptor, Cell-cell contact and Non-Protein signaling.

Pseudo-time trajectory analysis was conducted using the R package Monocle (v2.34). We use the new Cell Data Set function to model a negative binomial distribution. Size factors and dispersion were estimated using the estimate Size Factors and estimate Dispersions functions, respectively. Differential Gene Test function was used for differential expression analysis, and the reduced Dimension function was used for dimensionality reduction. Genes with q_val < 0.01 were selected, and cell trajectories were constructed using the set Ordering Filter function. To examine changes in pseudo-time trajectories of key genes during disease progression, cells were ordered along inferred trajectories from Control/Klk8^f/f^ to STZ/Klk8^f/f^ and from STZ/Klk8^f/f^ to STZ/Klk8^ΔEC^ using the order Cells function. Finally, the differential Gene Test function was used to identify genes that changed along the trajectory.

### Confirmation of KLK8 cleaving syndecan-4 (SDC4)

Human GECs were exposed to Ad-KLK8 for 72 hrs and the cells and supernatants were collected. The supernatants were concentrated by ultrafiltration (Millipore, Millipore, Bedford, MA), and were then separated by SDS-PAGE and transferred to a PVDF membrane. Measurement of SDC4 and KLK8 in the membrane was carried out by western blotting. In addition, the membrane was also stained with coomassie blue staining solution to identify cleaved bands. Two ∼15kDa bands were excised from the membrane, and N-terminal sequencing was then performed using the Edman degradation method at Biotech Pack Scientific (Beijing, China) on a PPSQ-33A system (Shimadzu, Kyoto, Japan).

Degradation of SDC4 directly by KLK8 was confirmed by incubation of recombinant His-tagged human KLK8 (rhKLK8; R&D systems, 2025-SE) with recombinant His-tagged human SDC4 (rhSDC4; R&D systems, 2918-SD-050). Briefly, equal volumes of diluted rhKLK8 and lsyl-endopeptidase (Wako BioProducts, 129-02541) in activation buffer (50 mM Tris, pH 8.0) were incubated at 37 °C for 30 min to activate rhKLK8. Increasing concentrations of activated rhKLK8 (1-100 ng/μl) diluted in assay buffer (50 mM Tris, pH 9.0) were incubated with rhSDC4 (10 ng/μl) at 37 °C for 1 hr and 5 hrs. Proteins were then harvested for measuring SDC4 by western blotting.

### Statistical Analysis

Statistical analyses were performed using SPSS 20 (SPSS Inc., Chicago, USA). All data are expressed as mean ± SEM. Normal distribution was assessed by Shapiro-Wilk test. Statistical significance was determined according to sample distribution and homogeneity of variance. Statistical comparisons between two groups were determined by two-tailed Student’s t test. One-way or two-way ANOVA with Bonferroni’s post hoc test was performed for comparisons among multiple groups. Regression analysis was used to determine correlations between variables. P < 0.05 was considered statistically significant.

### Other methods were provided in supplemental materials

#### Results

##### KLK8 is upregulated in glomerular endothelial cells (GECs) in DN and causes GEC dysfunction

To assess whether KLK family members are potentially involved in DN development, we first examined the mRNA expression levels of Klks in renal tissues of the mice with STZ treatment. Klk8 exhibited the highest induction in mRNA expression among Klks in renal tissues of the mice upon STZ treatment. Its protein levels were also increased in renal tissues of mice with STZ treatment (Figure 1A&B).

**Figure 1.**
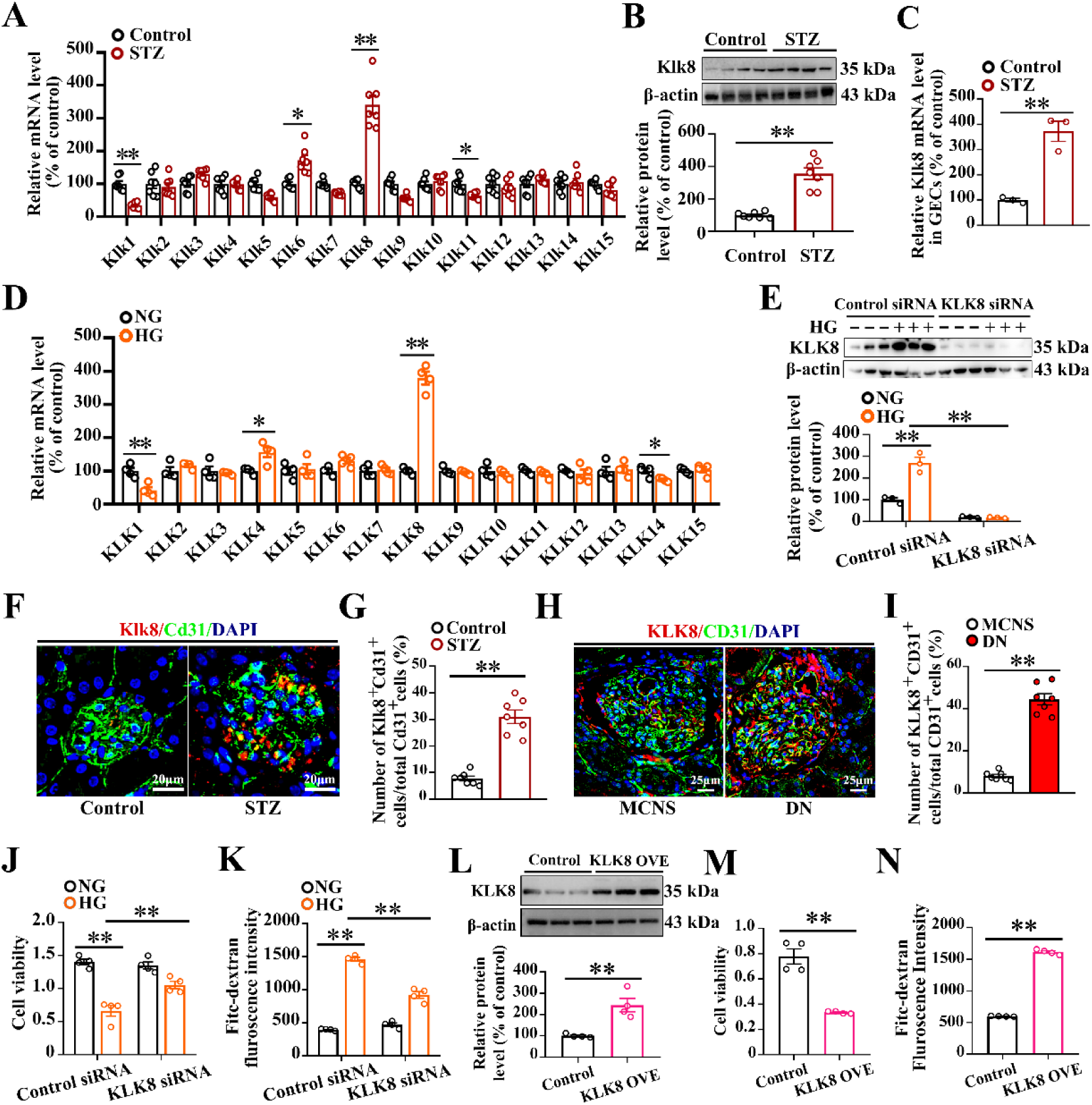
KLK8 expression is upregulated in GECs in diabetic mice and patients and involved in GECs dysfunction caused by HG. A-C, mouse diabetes was induced by STZ. The mRNA levels of Klks (A) and Klk8 protein level (B) in the renal tissues (n = 7). The mRNA level of Klk8 in GECs sorted by flow cytometry (C) (n = 3). D, human GECs were treated with HG for 120hrs, and the cells were harvested. The mRNA levels of KLKs in the cells (n = 4). E, human GECs were transfected with KLK8 siRNA and then treated with HG. The KLK8 protein level in the cells (n = 4). F&G, immunofluorescence analysis showed colocalization of Klk8 and Cd31 positive staining in glomeruli of diabetic mice. Nuclei were counterstained with DAPI (blue). The percentage of Cd31+/Klk8+ in the kidneys of control or diabetic mice was quantified (G) (n = 7). H&I, renal tissue biopsies were obtained from the patients with DN and MCNS. Immunofluorescence showed the colocalization of KLK8 and CD31 positive staining (H). Nuclei were counterstained with DAPI (blue). The percentage of CD31+/KLK8+ in the kidneys of patients with diabetes or MCNS was quantified (I) (n = 7). J&K, human GECs were transfected with KLK8 siRNA and then treated with HG. The cell viability (I) and permeability (J) were determined by CCK8 and FITC-dextran method, respectively (n = 4). L-N, GECs were infected with Ad-KLK8 and then KLK8 expression (L), cell viability (M) and permeability (N) were determined (n = 4). Data are expressed as means ± SEM. \**P* <0.05, ** *P* < 0.01.

GECs are emerging as a key player in DN pathogenesis and may initiate microalbuminuria, mesangial expansion, and podocyte injury in diabetes^20,21^. We then examined mRNA profile of KLKs in GECs upon HG and found that KLK8 mRNA level was also robustly induced and KLK8 protein expression was upregulated in cultured human GECs (Figure 1D&E). We sorted the GECs (Cd31^+^ and Cd45^-^) from the glomeruli using flow cytometry and found that Klk8 mRNA level in GECs of glomeruli increased by 3.72 ± 0.40-fold in STZ-treated mice compared to controls (Figure 1C). Using IF, we confirmed that glomerular Klk8 protein level in GECs was increased in STZ-treated mice compared to controls (Figure 1F&G). Moreover, a few KLK8 positive cells were detected in CD31+ cells in glomeruli of the patients with MCNS, but they were extensively increased in those of patients with diabetes (Figure 1H&I).

Next, we investigated whether increased KLK8 is involved in HG-induced GEC dysfunction. We confirmed that KLK8 siRNA caused a remarkable reduction of KLK8 expression (Figure 1E) and blocked the HG-induced decreased cell viability and prevented HG-induced GEC hyperpermeability (Figure 1J&K). Infection of GECs with adenovirus carrying KLK8 gene (Ad-KLK8) resulted in increased KLK8 expression, which caused reduced cell viability and increased endothelial permeability (Figure 1L-N).

### Global and endothelial-specific Klk8 deficiency attenuates hallmark features of DN in mice

To assess the roles of KLK8 in DN development, the Klk8^-/-^ mice and their littermates (Klk8^f/f^) were studied. The deletion efficiency of Klk8 was examined in renal tissues, Klk8 mRNA and protein levels were barely detectable in the kidneys of (Figure S2A&B). As expected, STZ caused a dramatic increase in serum glucose level (Figure S2C). The hallmark features of DN including reduced GFR and increased UACR and circulatory BUN level were found in Klk8^f/f^ mice at 16 weeks after STZ treatment compared to control Klk8^f/f^ mice (Figure 2A-D). Glomerular hypertrophy, mesangial matrix expansion, increased collagen, glycogen deposition and tubulointerstitial fibrosis were found in STZ-treated Klk8^f/f^ mice compared to control Klk8^f/f^ mice (Figure 2E). Glomerulosclerosis index and interstitial fibrosis were increased (Figure 2F&G), renal collagen I, hydroxyproline (HYP) and Tgf-β1 levels were increased in STZ-treated Klk8^f/f^ mice (Figure 2H-J). Of note, the above features of DN were attenuated in Klk8^-/-^ mice (Figure 2A-J). However, STZ-induced hyperglycemia was not alleviated in Klk8^-/-^ mice (Figure S2C). Transmission electron microscope (TEM) showed that loss of endothelial fenestration, thickened GBM and loss and fusion of podocyte foot processes in STZ-treated Klk8^f/f^ mice, which were improved in STZ-treated Klk8^-/-^ mice (Figure 2K).

**Figure 2.**
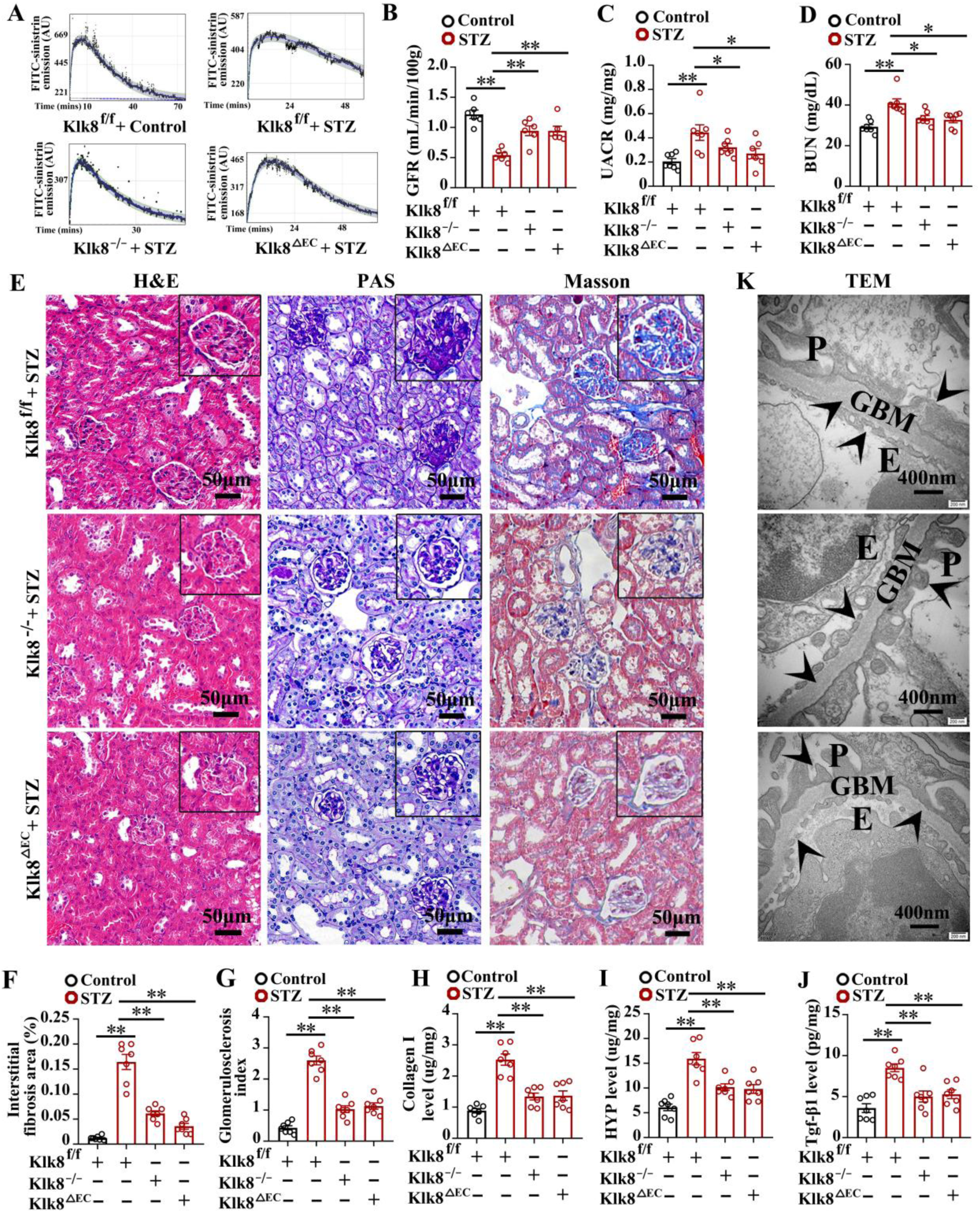
Global and endothelial Klk8 deficiency attenuates DN caused by STZ. Diabetes was induced by STZ in Klk8^f/f^, Klk8^-/-^ and Klk8^ΔEC^ mice. GFR via FITC-sinistrin clearance was determined in the mice at 16 weeks after STZ injection. A, the representative trace of GFR via FITC-sinistrin clearance. B, cumulative diagram of GFR clearance (n = 6). C&D UACR(C) and BUN level (D) in control and diabetic mice (n = 7). E, representative images of H&E, PAS and Masson’s trichrome staining (n = 7). F, quantification of interstitial fibrosis areas (n = 7). G, quantification of glomerulosclerosis index (n = 7). H, collagen-I level in renal tissues (n = 7). I, HYP level in renal tissues (n = 7). J, Tgf-β1 level in renal tissues (n = 7). K, representative images of TEM. Data are expressed as means ± SEM. * *P* < 0.05, ** *P* < 0.01. E, endothelial cells; P, podocytes; GBM, glomerular basement membrane.

Since KLK8 expression was extensively increased in GECs in DN, we study the roles of endothelial KLK8 in DN using Klk8^ΔEC^ mice. We confirmed that Klk8 expression was significantly decreased in GECs in Klk8^ΔEC^ mice compared to Klk8^f/f^ mice (Figure S3A). Reduced GFR, increased UACR and circulatory BUN level were significantly improved in STZ-treated Klk8^ΔEC^ mice compared to STZ-treated Klk8^f/f^ mice (Figure 2A-D). Glomerular hypertrophy, mesangial matrix expansion and collagen deposition and tubulointerstitial fibrosis caused by STZ were ameliorated in STZ-treated Klk8^ΔEC^ mice (Figure 2E). Glomerulosclerosis index, renal levels of collagen I, HYP, and Tgf-β1 were significantly reduced in STZ-treated Klk8^ΔEC^ mice compared to STZ-treated Klk8^f/f^ mice (Figure 2F-J). Loss of endothelial fenestration, thickened GBM and loss and fusion of podocyte foot processes caused by STZ were obviously reversed in Klk8^ΔEC^ mice (Figure 2K). Circulatory glucose levels were not differed between Klk8^ΔEC^ and Klk8^f/f^ mice after STZ treatment (Figure S3B).

Of note, there was no significant difference in GFR, UACR, BUN level and histology of renal tissues among Klk8^f/f^, Klk8^-/-^ and Klk8^ΔEC^ mice (Figure S4).

### Endothelial Klk8 regulates many pathways associated with diabetic kidney injury in many cells

To identify the molecular pathways changed in various cells linked to endothelial KLK8 in DN development, we performed scRNA-seq in diabetic Klk8^ΔEC^ mice. Analysis identified tubule cells including proximal tubule cells (PT), distal tubule cells (DT), collecting duct cells (DT) and loop of Henle cells (LOH), ECs, macrophages including kidney resident macrophages (KRM) and monocyte derived macrophages (Mo-Mφ), MCs, fibroblasts, podocytes and lymphocytes (T & NK cells and B cells) in kidney cortex (Figure 3A&B, supplemental File 1). As expected, Tie2 and Klk8 mRNA were mainly expressed in ECs and barely detectable in other cells (Figure S5A&B). Many pathways associated with DN in various cells in STZ-treated Klk8^f/f^ were reversed in STZ-treated Klk8^ΔEC^ mice. In ECs, 251 upregulated whilst 1239 downregulated GO-BP pathways in STZ-treated Klk8^f/f^ were reversed in STZ-treated Klk8^ΔEC^ mice, including extracellular matrix organization, response to stress, endothelial cell migration, regulation of apoptotic process involved in development, response to insulin, response to glucocorticoid (Figure 3C, supplemental File 2). In MCs, 829 pathways upregulated and 1002 downregulated by STZ in Klk8^f/f^ mice were reversed in Klk8^ΔEC^ mice, such as, extracellular matrix constituent secretion, smooth muscle cell proliferation, cell-substrate adhesion, extracellular matrix organization, regulation of basement membrane organization, mitochondrial electron transport, cytochrome c to oxygen. For PTC, 446 upregulated and 991 downregulated GO-BP pathways in STZ-treated Klk8^f/f^ mice were reversed in STZ-treated Klk8^ΔEC^ mice. These pathways included regulation of Wnt signaling, glucose metabolic process, cell surface receptor signaling pathway via JAK-STAT, chaperone-mediated autophagy, regulation of gap junction assembly. CellChat showed that the signaling pathways involved in crosstalk among many cells caused by STZ in Klk8^f/f^ mice were reversed in Klk8^ΔEC^ mice (Figure 3D). Of note, the pathways in ECs and MCs interacted other cells were increased in Klk8^f/f^ mice upon STZ treatment, which were remarkably reduced in Klk8^ΔEC^ mice.

**Figure 3.**
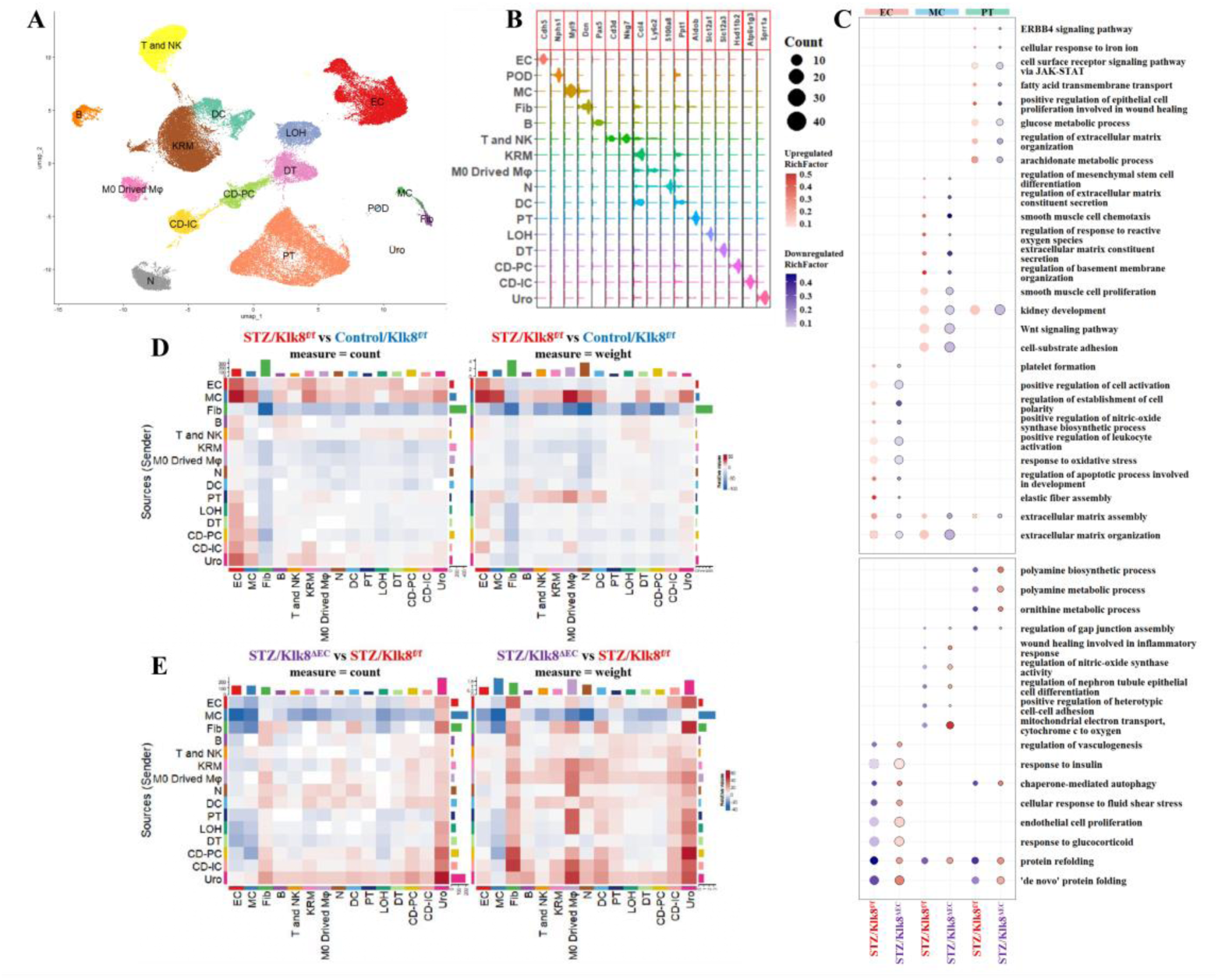
Endothelial Klk8 deficiency reverses the pathways related to DN in many cells revealed by scRNA-seq. Diabetes was induced by STZ in Klk8^f/f^ and Klk8^ΔEC^ mice. The renal tissues were obtained, and scRNA-seq was performed to determine transcriptome in various cells. A, scRNA-seq cell clusters in the diabetic kidney were visualized by uniform manifold approximation and projection (UMAP) after analysis using Seurat and R package. B, the markers of each cell cluster. C, the representative GO-BP pathways in ECs, MCs and PCs that were significantly changed in STZ/ Klk8^f/f^ mice upon vs Klk8^f/f^ mice and STZ/Klk8^ΔEC^mice with vs STZ/Klk8^f/f^ mice (P<0.05). D, cellchat showed the cell-communication among ECs, MCs, PCs, fibroblasts, macrophages, etc. between STZ-treated Klk8^f/f^ mice vs vehicle-treated Klk8^f/f^ mice. E, cellchat showed the communications among ECs, MCs, PCs, fibroblasts, macrophages, etc. between STZ/Klk8^ΔEC^ mice vs STZ/Klk8^f/f^ mice. KRM, kidney-resident macrophages; EC, endothelial cells; T and NK, T cells and NK cells; PT, proximal tubule; LOH, loop of henle; DT, distal tubule; N, neutrophils; CD-PC, principle cells of collecting duct; CD-IC, intercalated cells of collecting duct; M0 Drived Mφ, monocytes drived macrophages; B, B cells; DC, dendritic cells; Fib, fibroblasts; MC, mesangial cells; Uro, urotheliums; POD, podocytes.

### Endothelial KLK8 contributes to EC dysfunction by promoting endothelial-to-mesenchymal transition (EndMT) and SDC4 degradation in DN

We further analyzed the ECs and got 4 clusters, GECs, arterial ECs, capillary ECs and lymphatic ECs (Figure S6A, supplemental File.1). Given that GEC dysfunction is the early event in DN, we pay much attention to the changes in GECs. Of note, Klk8 expression was upregulated upon STZ in Klk8^f/f^ mice, which was blunted in Klk8^ΔEC^ mice (Figure S6B). Enrichment analysis showed the pathways associated with EndMT were increased in GECs of STZ-treated Klk8^f/f^ mice, in contrast, they were decreased in STZ-treated-Klk8^ΔEC^ mice (Figure 4A), such as extracellular matrix assembly, endothelial cell migration, smooth muscle cell migration, P53 signaling pathways, JAK-STAT signaling pathway, FoxO signaling pathway, AGE-RAGE signaling pathway in diabetic complications. CellChat showed that ECM-receptor pathways sent from GECs were extensively enhanced by STZ in Klk8^f/f^ mice, which were reversed in Klk8^ΔEC^ mice (Figure 4B&C). We then confirmed that EndMT took place in glomeruli in STZ-treated Klk8^f/f^ mice as evidenced by reduced Cd31 expression and increased vimentin and αSMA expression in endothelia of glomeruli compared to vehicle-treated Klk8^f/f^ mice (Figure 4D-F). These features were greatly improved in STZ-treated Klk8^ΔEC^ mice. Interestingly, our previous study showed that elevated KLK8 can cleave VE-cadherin, thereby leading to EndMT in ECs of heart^17^. In cultured human GECs, we found that KLK8 siRNA prevented reduced VE-cadherin level and increased αSMA and vimentin expression caused by HG (Figure 4G). We also confirmed that increased KLK8 could lead to EndMT in GECs by showing that infection of Ad-KLK8 resulted in an increase in KLK8 expression, which promoted αSMA and vimentin expression whilst suppressed VE-cadherin expression in GECs (Figure 4H). Figure 4I showed that elevated KLK8 expression extensively reduced VE-cadherin expression in GEC membranes.

**Figure 4.**
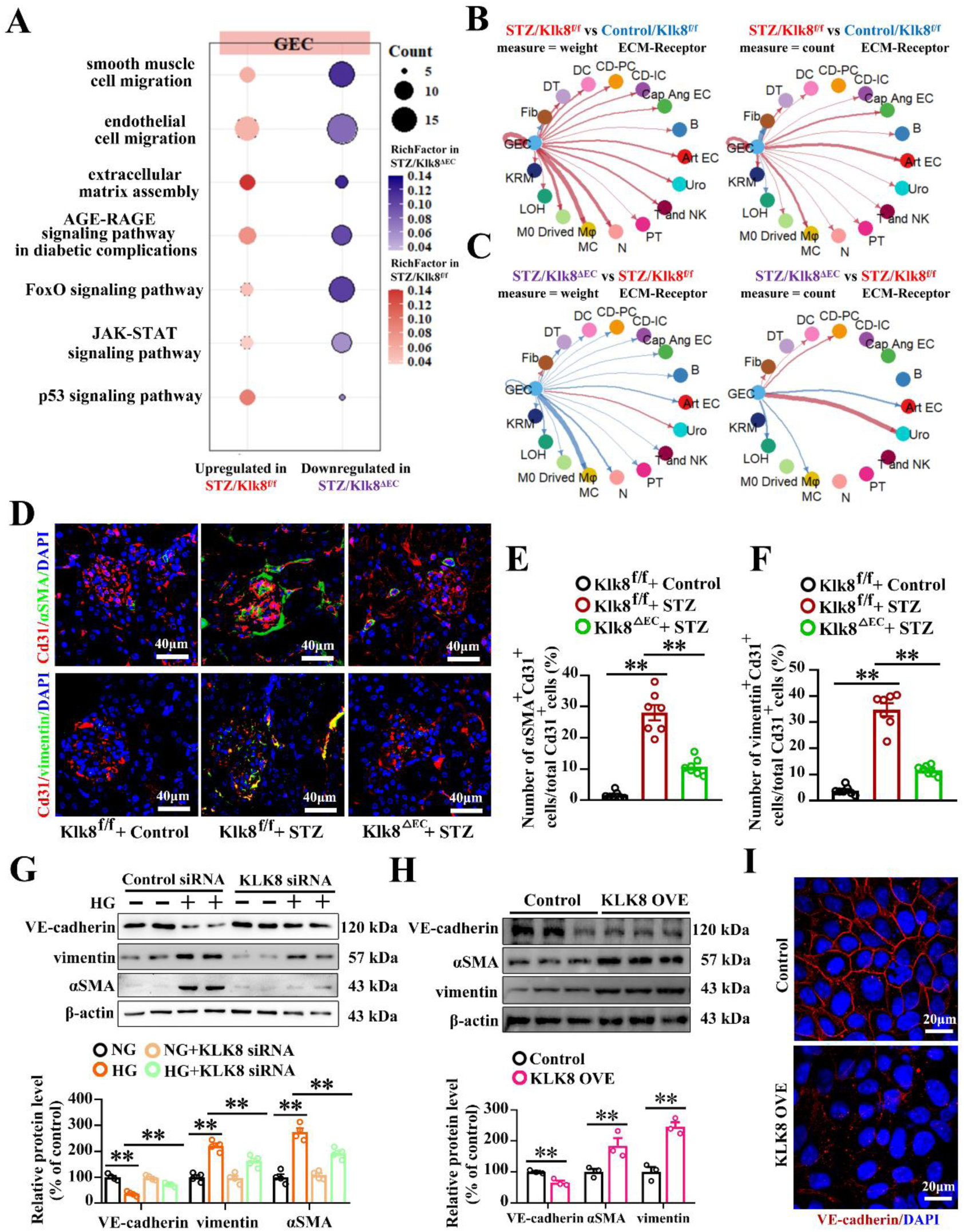
Endothelial Klk8 contributes to EndMT in DN. A, scRNA-seq showed that upregulated pathways associated with EndMT in GECs by STZ were reversed in Klk8^ΔEC^ mice. B&C, ECM-receptor pathways sent from GECs were enhanced by STZ(B), which were reversed in Klk8^ΔEC^ mice (C). D-F, EndMT took place in glomeruli in STZ-treated Klk8^f/f^ mice as showing that increased co-staining of αSMA with Cd31 (D&E) and vimentin with Cd31(D&F) in glomeruli of STZ-treated Klk8^f/f^ mice. Co-staining of αSMA with Cd31 (D&E) and vimentin with Cd31(D&F) was ameliorated in STZ-treated Klk8^ΔEC^ mice (n = 7). G, KLK8 siRNA prevented increased αSMA and vimentin expression and reduced VE-cadherin expression caused by HG treatment in cultured GECs (n = 4). H, Ad-KLK8 increased αSMA and vimentin expression and reduced VE-cadherin expression in GECs (n = 4). I, Ad-KLK8 impaired VE-cadherin staining in membrane in GECs. Data are expressed as means ± SEM. * *P* < 0.05, ** *P* < 0.01.

Next, we sought to identify potential proteins that can be degraded by KLK8 in human GECs by using Co-IP-MS. Among the potential targets, we noticed that SDC4 was identified in KLK8 Co-IP-MS (Table S5). Since SDC4 is a critical component of glycocalyx in endothelial cells, and SDC4 shedding is known to be a contributor to early DN^22^, we then focused on the cleavage effects of KLK8 on SDC4. We confirmed that SDC4 was co-immunoprecipitated by anti-KLK8 (Figure 5A). Incubation of rhSDC4 with rhKLK8 at 37°C caused a decrease of SDC4 in a time-dependent and dose-dependent manner (Figure 5B). In addition, we found that two new ∼15 kDa fragments of SDC4 were released into the culture media in GECs after transfection with Ad-KLK8 (Figure 5C). N-terminal sequencing was then performed on these fragments, and the data showed that KLK8 cleaved SDC4 before amino acid 115 in the extracellular region of SDC4, which provides convincing evidence that KLK8 directly cleaves SDC4 in GECs (Figure 5D&E).

**Figure 5.**
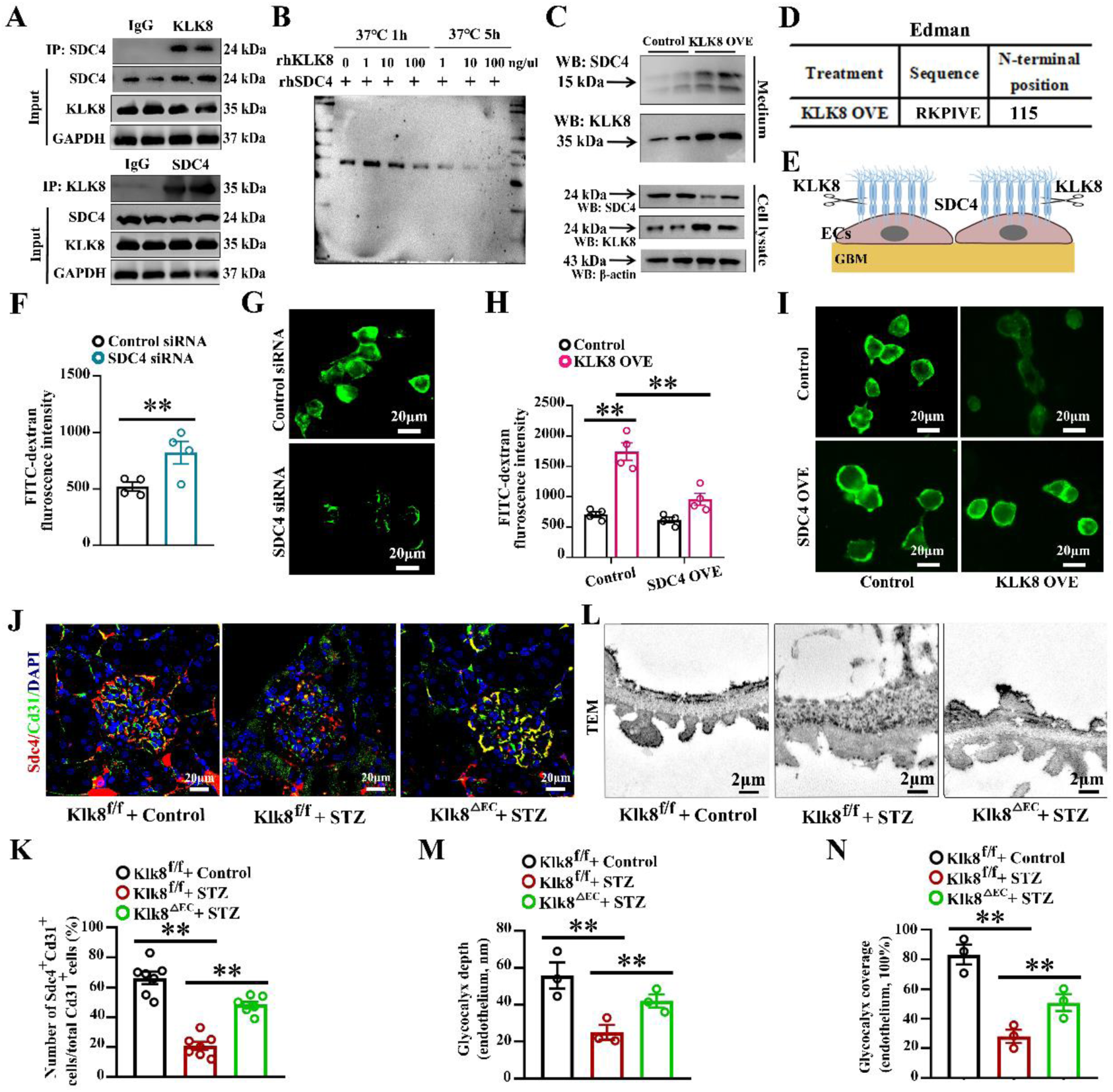
Endothelial Klk8 contributes to GEC dysfunction via cleavage of SDC4 in DN. A-E, KLK8 cleaved SDC4 in human GECs. A, binding of KLK8 to SDC4 was confirmed by reciprocal Co-IP in GECs. B, Incubation of rhSDC4 with rhKLK8 at 37°C for 1-5 hrs, and the levels of SDC4 was showed by western blotting. C, GECs were infected with Ad-KLK8 for 72 hrs. The cells and culture media were harvested for determination of SDC4 levels in cells and media. The upper panel showed two ∼15 kDa N-terminal SDC4 fragments in the media. D, the N-terminal sequence of two ∼15 kDa protein band was determined by Edman assay, and it showed that they belong to SDC4 sequence, and KLK8 cleaved SDC4 before amino-acid 115 in the extracellular domain. E, schematic diagram of the extracellular part of SDC4 cleaved by KLK8. F&G, reduced SDC4 caused dysfunction and damage in GECs. The GECs were transfected with SDC4 siRNA and control siRNA for 24 hrs. The cell permeability was determined by measuring FITC-dextran flux (F) (n = 4). FITC-WGA staining indicated that glycocalyx was damaged in the cells with SDC4 knockdown (G). H-I, increased permeability and glycocalyx damage caused by increased KLK8 expression could be reversed by SDC4 OVE in GECs. The GECs were infected with Lv-vector or Lv-SDC4. Then **t**he cells were then infected with Ad-vector or Ad-KLK8 for 72hr. The cell permeability was determined by measuring FITC-dextran flux (H) (n = 4). Glycocalyx was shown by FITC-WGA staining (I). J-N, diabetes was induced by STZ in Klk8^f/f^, Klk8^-/-^, Klk8^ΔEC^ mice. J&K, the Sdc4 expression in GECs in Klk8^f/f^ and Klk8^ΔEC^ mice with STZ treatment. J, representative image of Sdc4 in GECs in diabetic mice. K, the percentage of Cd31+/Sdc4+ in glomeruli in diabetic mice (n = 7). L, representative electron microscopy images of renal endothelial glycocalyx. M&N, quantification of the glycocalyx depth (M) and coverage (N) (n = 3). Data are expressed as means ± SEM. ** *P* < 0.01.

Next, we investigated whether loss of SDC4 confers KLK8-induced endothelial dysfunction in cultured human GECs. SDC4 siRNA resulted in about 80% reduction in SDC4 protein level (Figure S7A) and caused increased endothelial permeability (Figure 5F) and impaired endothelial glycocalyx by showing FITC-WGA (Figure 5G) and reduced cell viability (Figure S7B). Ad-KLK8 led to a significant decrease in FITC-WGA fluorescence intensity and increased cell permeability, which could be restored by Lv-SDC4 treatment (Figure 5H&I). Additionally, reduced cell viability and increased release of thrombomodulin, VWF and E-selectin caused by Ad-KLK8 were reversed by Lv-SDC4 treatment (Figure S7C-G).

In animal models, we found that Sdc4 expression level was significantly down-regulated in GECs of glomeruli in STZ-treated Klk8^f/f^ mice compared to vehicle-treated Klk8^f/f^ mice, and such alternation was significantly attenuated in Klk8^ΔEC^ mice (Figure 5J&K). Electron microscopy was then used to observe the ultrastructure of the endothelial glycocalyx stained with lanthanum nitrate. STZ treatment markedly decreased the endothelial glycocalyx coverage in the glomerulus. The glomerular endothelial glycocalyx loss by STZ in Klk8^f/f^ mice were greatly improved in Klk8^ΔEC^ mice (Figure 5L-N)

### Increased LIF/LIFR signaling contributes to EC dysfunction and mesangial cell activation linked to endothelial KLK8 in DN

Give that the secretory pathways received by GECs and MCs were increased in STZ-treated Klk8^f/f^ mice, which was decreased in Klk8^ΔEC^ mice (Figure S6C&D). We then used disease-related pseudo-time trajectories to identify the key genes of the receptors linked to Klk8 in DN development and found that Lifr was one of the key genes linked to Klk8 in ECs and MCs (Figure 6A&B). ScRNA-seq showed that Lifr mRNA was expressed in several cell types with high abundance in ECs and MCs (Figure S5C). We noticed that Lifr expression was markedly increased in both GECs and MCs in glomeruli upon STZ treatment in Klk8^f/f^ mice (Figure 6C-F). These alternations were substantially reduced in Klk8^ΔEC^ mice. In cultured GECs, we showed that HG caused LIFR upregulation, which was prevented by KLK8 siRNA (Figure 6G), while Ad-KLK8 enhanced LIFR expression (Figure 6H). We then treated the MCs with conditional media (CM) collected from GECs of Klk8 transgenic (Klk8 OVE) rats and found that it increased Lifr expression in MCs (Figure 6I). These data indicate that increased endothelial Klk8 contributes to upregulate Lifr expression in ECs and MCs during DN.

**Figure 6.**
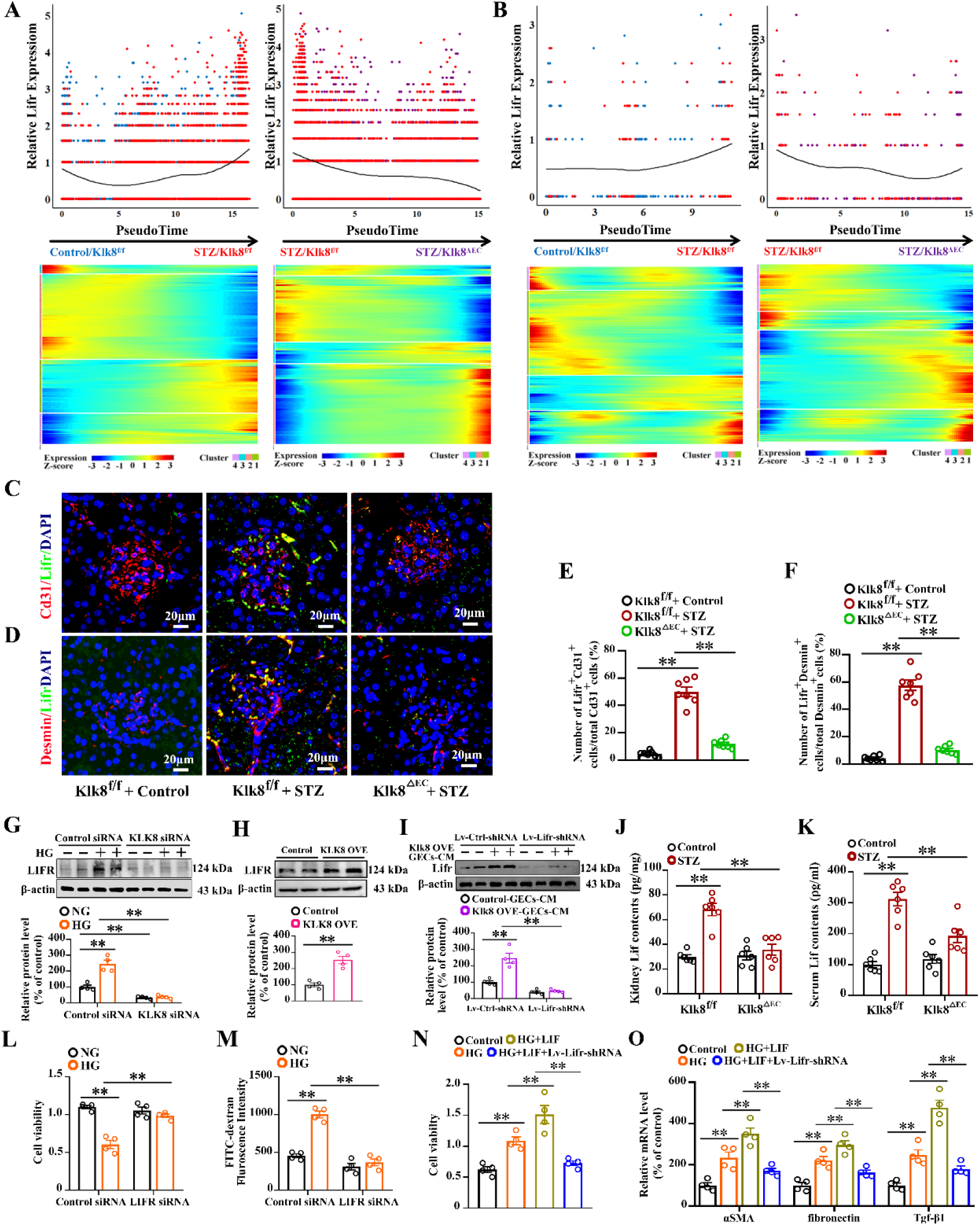
Increased LIFR signaling is involved in GEC dysfunction and MC activation linked to endothelial Klk8 during DN progress. A&B, pseudo-time trajectories showed Lifr expressions in ECs (A) and MCs (B) in STZ-treated Klk8^f/f^ and Klk8^ΔEC^ mice. C&E, increased Lifr expression in GECs caused by STZ in Klk8^f/f^ mice was reversed in Klk8^ΔEC^ mice. C, the representative images of co-staining of Lifr and Cd31 in glomeruli. E, the cumulative diagram of percentage of Lifr+/Cd31+ cells in glomeruli (n = 7). D&F, increased Lifr expression in MCs caused by STZ in Klk8^f/f^ mice was reversed in Klk8^ΔEC^ mice. D, the representative images of co-staining of Lifr and Desmin in glomeruli. F, the cumulative diagram of percentage of Lifr+/Desmin+ cells in glomeruli (n = 7). G&H, KLK8 modulates LIFR expression in cultured human GECs. Human GECs were transfected with KLK8 siRNA and then treated with HG (G) or GECs were infected with Ad-KLK8(H), and the cells were then harvested (n = 4). MCs were infected with Lv-Lifr shRNA and then treated with control-GECs-CM or Klk8 OVE GECs-CM (I) (n = 4). The Lifr expression was determined by western blotting. J&K, STZ increased Lif concentration in renal tissues (J) and blood (K) in Klk8^f/f^ mice, which was reversed by endothelial Klk8 deficiency (n = 7). L&M, LIFR knockdown by specific siRNA prevented HG-induced reduction of cell viability (L) and an increase in permeability (M) in GECs (n = 4). N&O, MCs were transfected with Lv-control shRNA or Lv-Lifr shRNA and then treated with LIF or HG. The cell viability was determined by CCK8 (N) and mRNA levels of αSMA, vimentin and Tgf-β1 were determined by Q-PCR (O) (n = 4). Data are expressed as means ± SEM. * *P* < 0.05, ** *P* < 0.01.

KLK8 is known to cleave the extracellular portion of several membrane proteins including PAR1 and PAR2, thereby leading to PAR1 and PAR2 activation^23,24^. Notably, the mRNA of F2r gene encoding PAR1 was found in ECs, in contrast, F2rl1 mRNA, which encodes PAR2, was barely detected in ECs (Figure S5D&E). We confirmed that the enhancement of LIFR expression by Ad-KLK8 was reversed by PAR1 antagonist (Figure S8A). The CM collected from Klk8 OVE GECs with PAR1 antagonist could not increase Lifr expression in MCs (Figure S8B). In addition, we found that LIF could increase KLK8 expression in GECs, which was reversed by LIFR siRNA (Figure S8C), indicating that reciprocal effects occur between KLK8 and LIFR signaling in ECs.

Ligand-receptor pairs analysis showed Osm-Lifr where ECs and MCs were the targeted cells whilst macrophages and neutrophils were the sources (Figure S6E). In fact, LIF is the classic ligand of LIFR and is produced by many cells including ECs, immune cells, etc^25–27^. It is necessary to examine the Osm and Lif protein levels in circulation and kidney tissues in response to STZ. We found that there was no obvious change in Osm levels in plasma and kidney tissues upon STZ treatment in Klk8^f/f^ mice (Figure S9). In contrast, Lif levels in plasma and renal tissues were significantly increased in STZ group compared to controls. Interestingly, its levels were reduced in STZ-treated Klk8^ΔEC^ mice compared to STZ-treated Klk8^f/f^ mice (Figure 6J&K).

Next, we investigated the roles of LIFR signaling in HG-induced GEC dysfunction and MC activation in cultured cell models. By using LIF treatment to activate LIFR, we found that LIF treatment increased cell permeability and reduced cell viability in GECs, which was prevented by LIFR siRNA (Figure S10). Reduced cell viability caused by HG was reversed by LIFR siRNA in GECs (Figure 6L&M). LIF significantly promoted HG-induced proliferation and activation markers in MCs as evidenced by enhanced cell viability and upregulated expression of fibronectin, Tgf-β1 and αSMA, which was reversed by Lifr knockdown (Figure 6N&O).

CellChat analysis showed that interaction between GECs and MCs were extensively increased in Klk8^f/f^ mice after STZ treatment, which was decreased in Klk8^ΔEC^ mice (Figure 3D&E). Particularly, the enhanced secretory pathways sent from GECs to MCs caused by STZ were reduced in Klk8^ΔEC^ mice (Figure S6C&D), indicating that Klk8 enhances secretory pathways sent from GECs. In cultured MCs, we found that CM of GECs with Klk8 OVE significantly increased cell proliferation and increased fibronectin, Tgf-β1 and αSMA expression compared to CM of control GECs (Figure S11A-D). However, the CM from the Klk8 OVE GECs with PAR1 antagonist treatment could not increase cell viability, fibronectin Tgf-β1 and αSMA expression (Figure S11A-D). Using secretome combined with ELISA, we identified that LIF level was increased in the supernatants of GECs with Klk8 OVE, which was reversed by PAR1 antagonist (Figure S11E-G). Both LIF neutralizing antibody and Lifr shRNA reversed the effects of CM of Klk8 OVE GECs on proliferation and activation MCs (Figure S11H-O).

### Lv-Lifr shRNA ameliorates hallmark features of DN in mice

Next, we elucidate the roles of LIFR signaling in the progression of STZ-induced DN. Given that Lifr is expressed in ECs and MCs in kidney, we constructed a lentiviral vector incorporating Lifr-shRNA to achieve comprehensive reduction of Lifr expression in renal tissues. As expected, Lifr level was robustly increased in renal tissues upon STZ treatment, which was blunted by Lv-Lifr shRNA treatment (Figure 7A). Increased Lifr expression in ECs and MCs caused by STZ was reversed by Lv-Lifr shRNA treatment (Figure 7B-D).

**Figure 7.**
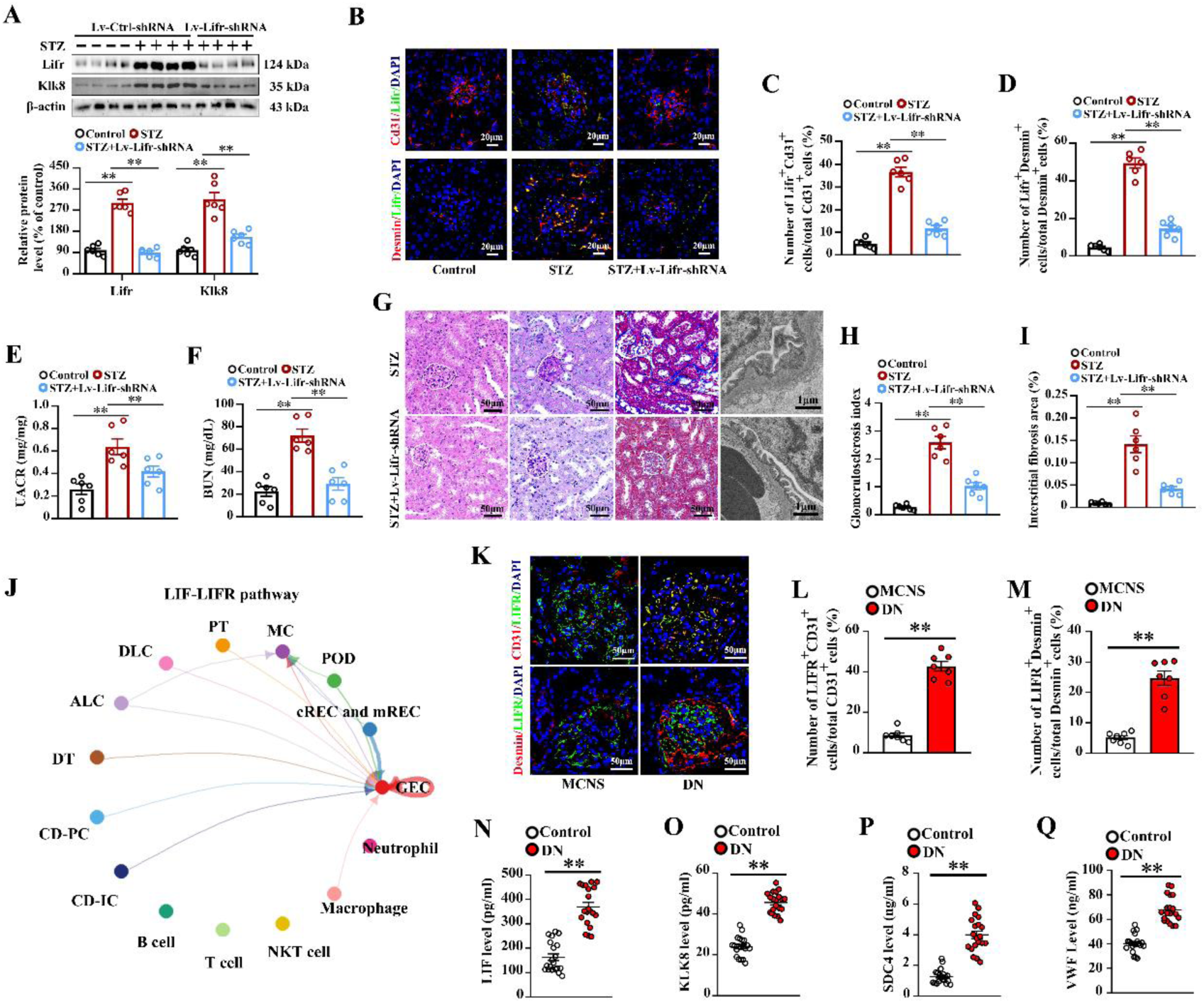
Lifr knockdown ameliorates features of DN in mice and circulatory KLK8 level is positively correlated with sSDC4 in patients with diabetes. A-I, diabetic mice was induced by STZ. The mice were then injected with Lv-Lifr shRNA (i.v) at 7^th^ day after STZ treatment. The features of DN were determined at 16^th^ week after STZ treatment. A, Lifr and Klk8 levels in renal tissues were determined by western blotting (n = 6). B-D, Lifr expression in GECs and MCs in glomeruli of the diabetic mice (n = 6). E&F, UACR (E) and BUN level (F) in diabetic mice (n = 6). G, representative images of H&E, PAS, Masson’s trichrome staining and TEM. H, quantification of glomerulosclerosis index (n = 6). I, quantification of interstitial fibrosis areas (n = 6). J, CellChat showed LIF-LIFR communication among GECs, MCs, PCs, podocytes and immune cells in renal tissues of the patients with diabetes. K-M, LIFR expression in GECs and MCs in glomeruli of diabetic patients (n = 7). N-Q, circulatory levels of LIF (N), KLK8 (O), soluble SDC4 (P) and VWF (Q) in patients with diabetes (n = 20). Data are expressed as means ± SEM. ** *P* < 0.01.

Increased UACR and circulatory BUN level, glomerular hypertrophy, collagen deposition and tubulointerstitial fibrosis in diabetic mice were significantly improved after Lv-Lifr-shRNA treatment (Figure 7E-I). Loss of endothelial fenestration, thickened GBM and podocyte foot process loss and fusion in STZ-treated mice were obviously attenuated by Lv-Lifr shRNA (Figure 7G). Reduced SDC4 expression in GECs were reversed by Lv-Lifr shRNA in diabetic mice (Figure S12). In addition, EndMT in glomeruli of diabetic mice was reversed by Lv-Lifr shRNA as evidenced by increased Cd31 expression, and decreased αSMA and vimentin expression in mice with Lv-Lifr shRNA compared with controls (Figure S13). In addition, we found that the upregulated Klk8 expression in renal tissues caused by STZ was reversed by Lv-Lifr shRNA (Figure 7A).

### Circulatory LIF and renal LIFR expression are increased and circulatory KLK8 level is positively correlated with LIF, VWF and creatinine levels in patients with DN

We subsequently analyzed scRNA-seq data from the patients with DN (GSE183279) and found that LIF-LIFR was one of the ligand-receptor pairs where GECs were the targeted cells in the patients with DN (Figure 7J). We then confirmed that LIFR expression was significantly increased in GECs and MCs in DN patients compared to MCNS patients (Figure 7K-M). Interestingly, the levels of LIF, KLK8, soluble SDC4 and endothelial injury biomarker VWF in circulation were increased in patients with DN compared to healthy volunteers (Figure 7N-Q). Correlation analysis showed that KLK8 level was positively correlated with levels of LIF and VWF as well as creatinine in DN patients (Figure S14).

## Discussion

Some studies have demonstrated that several endothelial factors such as VEGF, eNOS and LRG1 play important roles in DN pathogenesis^3–5^; however, the key pathways governing EC dysfunction in DN progress remain poorly understood. The present study has provided novel findings regarding the importance of endothelial KLK8 in DN pathogenesis. First, we demonstrated that not only global but also endothelial Klk8 knockout prevented hallmark features of DN including albuminuria, reduced GFR and glomeruli sclerosis and interstitial fibrosis caused by STZ in mice. Second, we revealed that endothelial KLK8 was at least involved in two important events EndMT and impairment of glycocalyx integrity in GECs in DN. By using proteomic analysis coupled with biochemical experiments, we revealed that KLK8 directly cleavage SDC4, thereby leading to loss of glycocalyx integrity of GECs. Third, scRNA-seq analyses revealed the main pathways related to DN in endothelial cells, MCs, and TCs were reversed by endothelial Klk8 deficiency. Furthermore, we identified that Lifr was one of key genes regulated by endothelial Klk8 during DN, and Lifr knockdown ameliorates hallmark features of DN in mice.

As mentioned, KLKs exert their function via cleaving peptide bonds of targeted proteins. Our study identified a novel targeted protein of KLK8-SDC4. We demonstrated that KLK8 could act as a “sheddase” to cleave the extracellular region of SDC4 from GECs, thereby contributing to loss of glycocalyx integrity in GECs in DN. Impairment of glycocalyx integrity of GECs is believed to be associated with the development of microalbuminuria in type 1 and type 2 diabetes^28,29^. First, we demonstrated that glycocalyx in GECs was intensively decreased upon STZ treatment, which was reversed by endothelial Klk8 deficiency. The endothelial glycocalyx is composed chiefly of proteoglycans, which have a core protein like SDCs and glycosaminoglycan side chains like heparan sulfate (HS). The SDCs can be cleaved from the cell surface by “sheddases” including matrix metalloproteinases and a disintegrin and metalloproteinase-17^22,30^, thereby leading to glycocalyx loss. Second, using biochemical approaches, we provided convincing evidence that KLK8 cleaves the extracellular region of SDC4 from GECs. Finaly, we confirmed that SDC4 mediated KLK8 induced glycocalyx integrity loss and dysfunction in GECs in cell model. Thus, our study strongly indicates that endothelial KLK8 contributes to GEC dysfunction via cleavage of SDC4.

Our previous study has identified that VE-cadherin is a substrate of KLK8 in ECs. We have demonstrated that KLK8 degrades VE-cadherin in ECs of heart, which causes plakoglobin nuclear translocation and subsequently activates p53 signaling pathway, leading to EndMT in diabetic cardiopathy^17^. In fact, EndMT is a distinguishing event during DN progress^31^. In consistence with the findings in cardiac ECs, we demonstrated that endothelial-specific KLK8 deficiency mitigated EndMT in glomeruli, restored VE-cadherin expression, and suppressed the upregulated p53 signaling pathway in GECs during DN. Conversely, increased KLK8 levels promoted EndMT in cultured GECs, suggesting that similar mechanisms underlie KLK8-induced EndMT in both cardiac and renal ECs.

The scRNA-seq analysis indicates that endothelial Klk8 is involved in many pathways associated with kidney injury of DN in many cells such as MCs and DTs, which supports the theory that ECs are the key players in DN progress. Moreover, our data indicate that endothelial KLK8 is involved in abnormal communication of ECs with MCs in DN. The MCs are centrally located in the capillary tuft, between the glomerular capillaries, thereby providing key structural support for glomerular capillary loops and mediating glomerular crosstalk. In DN, MCs become activated resulting in mesangial expansion that occurs before the onset of clinical manifestations. Emerging evidence indicates that MCs play central roles in DN development^32^. Liu et al. have demonstrated that the pathways related to MC activation, such as extracellular matrix organization, regulation of cell adhesion, regulation of smooth muscle proliferation and positive regulation of cell motility in DN^33^. Consistently, these pathways were also found in the present study. Moreover, we revealed that these pathways were related to endothelial Klk8 in DN. The MCs are in direct contact with GECs. Cellchat analysis revealed that enhanced secretory pathways sent from GECs to MCs were reversed by endothelial Klk8 deficiency. In the cell model, we showed that the CM of Klk8 OVE GECs led to activation of MCs, which provides evidence that KLK8 in GECs can activate MCs via the secretory mediators of ECs. Moreover, we found that LIF would be one of the secretory mediators in ECs induced by KLK8 for MC activation.

Some studies have demonstrated the involvement of LIFR signaling in renal interstitial fibrosis. Upregulation of Lifr in renal tissues are found upon ischemia-reperfusion and unilateral ureteral obstruction^25,34,35^. LIFR signaling contributes to renal fibrosis progress by promoting EMT in tubular cells^34^ and crosstalk between fibroblast and tubular cells^25^. To our knowledge, the roles of LIFR in DN have not been reported. In the present study, we revealed that LIFR signaling mediated many functions of endothelial KLK8 in DN progress. First, pseudotime trajectory demonstrated dynamic changes of Lifr expression in ECs and MCs as the disease progressed, which were reversed by endothelial Klk8 deficiency. Then, using cell model, we revealed that the upregulated LIFR was also involved in GEC dysfunction caused by HG. Moreover, we confirmed that LIFR was involved in HG-induced MC activation, which was associated with endothelial KLK8. Endothelial KLK8 activated PAR1, thereby leading to upregulation of LIFR in ECs and MCs. Furthermore, we showed that Lifr knockdown by Lv-Lifr shRNA ameliorated the features of DN caused by STZ including albuminuria, mesangial expansion, glomerulosclerosis and interstitial fibrosis in mice. In addition, we found that increased Klk8 expression was also associated with increased Lifr in animal models and cell models, which indicates that feed-forward loop may exist between KLK8 and LIFR in DN progress. Thus, the effects of LIFR signaling on kidney injury in DN might be associated with KLK8.

## Conclusion

The present study revealed a novel mechanism underlying the pathogenesis of DN, specifically, the key roles of endothelial KLK8 in DN development. Elevated endothelial KLK8 contributes to GECs dysfunction by leading to loss of glycocalyx integrity by directly degrading SDC4 and promoting EndMT by cleaving VE-cadherin, and upregulation of LIFR signaling via PAR1 activation in DN. Endothelial KLK8 contributes to DN development by dominating abnormal communication with MCs through a LIFR-dependent mechanism (Graphical abstract). Our study identifies a novel driver of DN development and progress and immediately highlights potential therapeutic strategies targeting KLK8/LIFR signaling pathway for DN.

## Author contribution

X.N. conceived, designed, supervised the study, and wrote the manuscript. X.Z. supervised the study and wrote some parts of the manuscript. J.D carried out animal experiments, some in vitro cell experiments, manuscript preparation, IHC analysis and statistical analysis; M.L, carried out all the bioinformatic analysis and some data processing; Y.J carried out in vitro cell experiments and IHC and IF analysis. Z.T. carried out the genotype of genetically modified animals and animal experiments. D.X, J.Y, H.Y, S.P. provided clinical specimens and supervised the enrolment of DN patients; D.X analyzed morphology of renal tissues.

## Funding

This study was supported by National Natural Science Foundation of China (No. 82000797&82471711), Human Provincial Science and Technology (2018RS3030) and Hunan Provincial Natural Science Foundation (2021JJ31058).

## Availability of data and material

All data associated with this study are available from the corresponding author with reasonable request.

## Declarations

### Ethics approval and consent to participate

The collection of human kidney samples was carried out in accordance with ethical approval from the Medical Research of Xiangya Hospital Central South University (No.2023091177). Approval for all animal experiments was granted by the Laboratory Animal Ethics Committee of Navy Medical University.

### Consent for publication

All authors have approved the submission of this manuscript to this journal.

### Competing interests

The authors declare that there is no confict of interest

## Notes

### Competing Interest Statement

The authors have declared no competing interest.

